# The Elg1 Replication Factor C-like complex safeguards cells from replication stress through a noncanonical pathway independent of the Mec1-Rad53 axis

**DOI:** 10.1101/2025.05.29.656775

**Authors:** Pallavi Bose, Soumitra Sau

## Abstract

The Elg1 Replication Factor C-like complex (Elg1-RLC) that functions as a PCNA unloader, is known to be involved in multiple DNA replication/repair-related activities from yeast to humans. By exploiting disassembly-prone PCNA mutants, we reveal that Elg1-RLC uses its PCNA unloading activity to counter the DNA-alkylating agent methyl-methanesulfonate (MMS)-mediated slow progression of replication forks. Despite having a largely functional DNA Damage Response (DDR), the viability loss of *elg1Δ*-*DDR* double mutants, in the presence of MMS, matches that of *mec1Δ* and *rad53Δ* cells, deficient for the central checkpoint kinases. This suggests that *elg1Δ*-*DDR* double mutants experience replication fork collapse when exposed to MMS. Indeed, in response to MMS, accumulation of Rad52 foci in the replicative *elg1Δ*-*DDR* cells supports this possibility. However, the failure of rescuing *elg1Δ*-*DDR* mutants by elevating dNTP levels (by deleting the ribonucleotide reductase *SML1*) eliminates the possibility of a Rad53-regulated dNTP shortage-mediated fork collapse. Thus, we propose a S-phase checkpoint regulatory role of Elg1-RLC that works through a noncanonical pathway parallel to the canonical one. Collectively, our findings suggest a model in which Elg1-RLC, by timely unloading chromatin-bound PCNA from the damaged/stalled forks, coordinates the DDR pathways to safeguard the integrity of replication forks under replication stress.

## Introduction

Cell viability and proliferation depends on an accurate DNA copying mechanism followed by equal segregation of the replicated chromosomes into mother and daughter cells. In budding yeast, DNA replication is carried out by the replisome, a protein complex composed of many proteins including strand-specific DNA replicative polymerases and their processivity factor PCNA (Proliferating Cell Nuclear Antigen) (1). PCNA, a doughnut-shaped homotrimeric protein, encircles the double-stranded DNA and acts as a replication fork-associated sliding clamp (1–3). Apart from providing the physical support to DNA polymerases during replication, PCNA is involved in many other DNA/chromatin transactions processes: regulation of DNA replication, DNA repair, chromatin assembly and remodeling, and sister chromatid cohesion (1,4,5). These functions are executed by various PIP (PCNA Interacting Peptide)-motif containing proteins which are recruited at the functional sites by interacting either with the unmodified or modified (ubiquitylated or SUMOylated) PCNA (1,4). After completion of the Cdc9-mediated ligation of the terminal nucleotides of each ORI-generated replicating fragment, PCNA is unloaded by the Elg1 complex, known as Elg1-RLC (Elg1-Replicator Factor C-like complex), composed of Elg1 and the Rfc2-5 subunits (6). Three independent yeast genetic screenings identified Elg1 as a DNA replication protein and a regulator of genome-wide recombination events (7–9). Later, genetic and biochemical experiments established Elg1-RLC as a primary unloader of unmodified as well as modified PCNA from yeast chromatin (10–12). Due to their inability to unload PCNA from DNA, *elg1* mutants accumulate chromatin-bound PCNA and SUMOylated PCNA, which mediate genomic instability phenotypes such as increased sister chromatid recombination (13), sensitivity towards genotoxic agents (12,13), elongated telomeres (13,14), sister chromatid cohesion defects (15), defective nucleosome organization (16), inefficient chromatin silencing (17) and compromised mismatch repair (18). The Elg1-RLC is evolutionarily conserved, and its mammalian ortholog is called ATAD5 (hELG1) (19). Structurally and functionally hELG1 is analogous to yeast Elg1, and unloads ubiquitinated as well as unmodified PCNA from human chromatin (20,21). Absence of hELG1/ATAD5 causes cancer development in mice and humans (22). hElg1 is a component of the Fanconi Anemia (FA) pathway, and mutations in ATAD5 were found to be linked with different types of human cancer (22–24).

During DNA replication, cells may encounter various genomic insults that could result either in DNA damage-mediated slowing down of the replication fork (RF) progression or the complete stalling of RFs (25,26). Anomalies in RF movement or structure generate replication stress, and to cope up, cells activate the S-phase checkpoint (also known as DNA Damage Response or DDR), which functions through two sub-pathways: the DNA Damage Checkpoint (DDC), which is primarily triggered by the damaged DNA that impedes RF progression, and the DNA Replication Checkpoint (DRC), activated by arrested RFs (25–27). The essential function of the S-phase checkpoint is to maintain the viability of the replication-stressed cells by protecting the integrity of their replication forks, and an activated Mec1 – Rad53 axis executes this function (28,29). Both checkpoint branches initiate the signaling process by recruiting the apical kinase Mec1 (ATR in humans) and its co-factor Ddc2 (ATRIP in humans) to the RPA-coated single-stranded DNA regions (which act as stress signals) that result from either damaged DNA or stalled RFs (30,31). The two branches converge in activating the checkpoint effector kinase Rad53 (CHK1 in humans) (25). In parallel to Mec1-Ddc2 recruitment, a PCNA-like heterotrimeric clamp complex composed of Ddc1, Mec3 and Rad17 (the 9-1-1 complex) is recruited at the stressed chromatin sites by Rad24 (RAD17 in humans), the largest subunit of the Rad24-RLC (32,33). Thus, Rad24 and the 9-1-1 complex together act as sensor proteins for both the S-phase checkpoint branches (25,32). In the DDC branch, Rad9 (53BP1 in humans) acts as a mediator protein that assembles at the damaged sites through phosphorylated histone H2A and methylated histone H3, and becomes fully phosphorylated by Ddc1-activated Mec1 (34,35). Rad53 is recruited to the damaged chromatin sites by interacting with the phosphorylated Rad9 (34,36), following which Mec1 phosphorylates Rad53, which in turn undergoes autophosphorylation and becomes fully active (37). In the DRC branch, the replisome component Mrc1 (CLASPIN in humans) acts as a mediator protein, which is phosphorylated by Mec1 upon RF stalling (26,38). Once phosphorylated, Mrc1 along with Ctf18 (CTF18 in humans) of the Ctf18-RLC and the Sgs1 DNA helicase (BLM and WRN in humans) recruit Rad53 to the stalled fork (25). Mec1 phosphorylates Rad53 and stimulates its autophosphorylation (25). By phosphorylating the downstream effector checkpoint substrates Dun1 and Sml1, activated Rad53 stabilizes stalled RFs, upregulates the nucleotide pool, represses late origin firing and arrests the cell-cycle progression (25,39). Although Rad9 and Mrc1 define the DDC and DRC branches of the S-phase checkpoint, respectively, in response to stressed forks both proteins cooperate with each other and control the kinetics of Rad53 activation for an optimal completion of DNA replication under stress conditions (26).

Although several lines of evidence have emerged from previously published reports suggesting that the Elg1-RLC may play a role in the DDR pathway (8,40,41), the precise function remained unclear. In this study, we thought to understand the S-phase checkpoint-specific role of Elg1-RLC by combining a deletion of *ELG1* with DDR pathway mutants. Even after retaining almost a fully active Mec1-Rad53 axis, *elg1Δ*-*DDR* double mutants display higher MMS-sensitivity compared to their corresponding single mutants and experience cell death similarly to the central checkpoint mutants *mec1Δ/rad53Δ*. Thus, it appears that Elg1-RLC and DDR components are working in concert to protect the integrity of replication forks during replication stress through a Rad53-independent pathway.

## Results

### *ELG1* shows synthetic genetic interactions with DNA Damage checkpoint components in response to MMS-induced replication stress

Previous work has shown that Elg1 undergoes Mec1-dependent phosphorylation in response to MMS exposure (40). Moreover, in an artificially induced DNA Damage Checkpoint (DDC) mimicking system it was shown that Elg1 is required to phosphorylate Rad53 (41). These results point to a possible involvement of Elg1 in the DDR. Since both *elg1Δ* cells and *DDR* mutants show sensitivity to MMS (Fig. S1A), we tested the relationship between Elg1 and the DDR by creating a collection of *elg1Δ-DDR* double mutants, in which different participants of the DDR were deleted in the *elg1Δ* background, and their growth on MMS-containing plates was compared to that of corresponding single mutants.

Under replication stress, DDR sensors are the first set of proteins of the DDR checkpoint pathway to be recruited at the stressed sites marked by RPA-coated single stranded DNA (33). First, we deleted DDR sensors: the members of the 9-1-1 complex (*DDC1*-*MEC3-RAD17* in yeast; RAD9-RAD1-HUS1 in humans) and their loader *RAD24* in the *elg1Δ* background, and the generated double mutants were spotted on plates supplemented with MMS. MMS-mediated DNA methylation slows down the progression of replication forks resulting replication stress, and hence triggers the S-phase checkpoint (42). *elg1Δ* cells and single *DDR* sensor mutants (*ddc1Δ*, *mec3Δ*, *rad17Δ* and *rad24Δ*) failed to grow at 0.012% and 0.015% MMS, respectively. The double mutants (*elg1Δ ddc1Δ*, *elg1Δ mec3Δ*, *elg1Δ rad17Δ* and *elg1Δ rad24Δ*) exhibited a synergistic phenotype: they were ∼8 to 10-fold more sensitive to MMS compared to the corresponding single mutants (Fig. 1, A-D).

**Figure 1.**
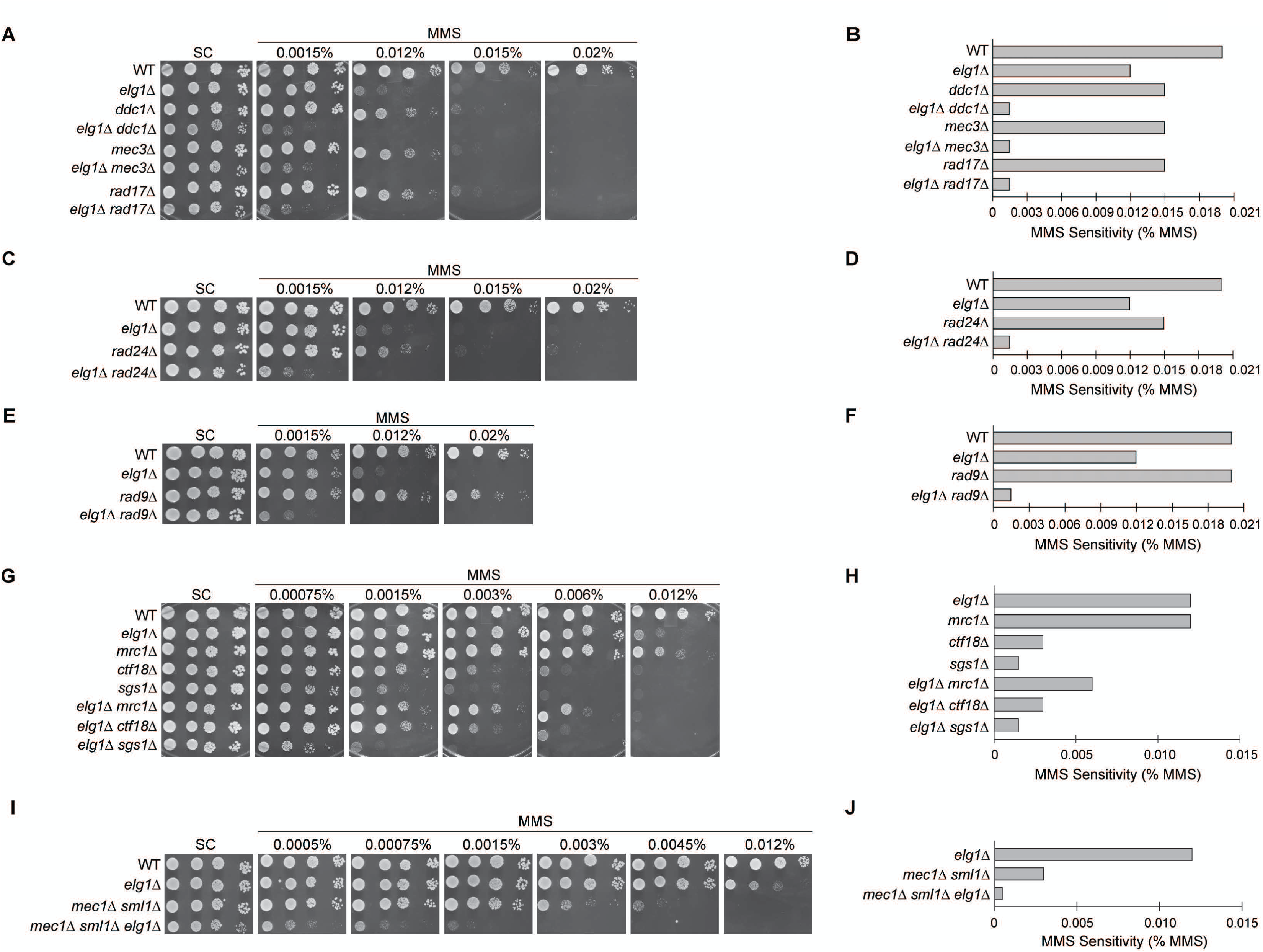
*ELG1* and DNA Damage checkpoint components exhibit synergistic interactions in response to MMS. Ten-fold serially diluted overnight cultures of wild-type (WT) and indicated single, double and triple mutants were dropped on regular synthetic complete (SC) plates and SC plates supplemented with indicated amounts of MMS. The MMS sensitivity of each strain was recorded from the drop assay plates and displayed as a bar diagram. (A – D) Spot assays and MMS sensitivity index for *elg1Δ*-*DDR* sensor double mutants (*elg1Δ ddc1Δ*, *elg1Δ mec3Δ*, *elg1Δ rad17Δ* and *elg1Δ rad24Δ*) and their corresponding single mutants. (E – F) Spot assays and MMS sensitivity index for *elg1Δ*-*DDC* mediator double mutant (*elg1Δ rad9Δ*) and its corresponding single mutants. (G – H) Spot assays and MMS sensitivity index for *elg1Δ*-*DRC* mediator double mutants (*elg1Δ mrc1Δ, elg1Δ ctf18Δ* and *elg1Δ sgs1Δ*) and their corresponding single mutants. (I – J) Spot assays and MMS sensitivity index for *mec1Δ sml1Δ elg1Δ* triple mutant and its corresponding single and double mutant. In each plate, WT spots were used as a control. The drop assay for each strain was repeated at least three times (including WT and *elg1Δ*) with three independent transformants (except WT and *elg1Δ*), however, for the representation purpose, images from only one set is provided here. For Figures 1, 2 and 4F: all SC plates ± MMS were kept at 28°C. Images of SC plates – MMS were recorded after 2 days of incubation, whereas SC plates + MMS were recorded after 3 days of incubation.

After finding that *ELG1* and the DDR sensors interact synergistically, we created and tested the growth of *elg1Δ*-*DDC* mediator double mutant (*elg1Δ rad9Δ*) and *elg1Δ*-*DRC* mediator double mutants (*elg1Δ mrc1Δ, elg1Δ ctf18Δ* and *elg1Δ sgs1Δ*) on MMS-containing plates. Similar to the *elg1Δ*-*DDR* sensor double mutants, the *elg1Δ*-*DDC* mediator double mutant (*elg1Δ rad9Δ*) displayed a synergistic phenotype: it was 8– and ∼13-fold more sensitive to MMS compared to *elg1Δ* and *rad9Δ* cells, respectively (Fig. 1, E and F). Cells lacking Mrc1 showed mild sensitivity to 0.012% MMS, whereas the *elg1Δ*-*DRC* mediator double mutant *elg1Δ mrc1Δ* displayed a *rad53Δ sml1Δ*-like defective growth pattern on synthetic complete plates (data not shown) and was found to be more sensitive to MMS compared to its corresponding single mutants (Fig. 1, G and H). This suggests that Elg1 and Mrc1 together play a role to upkeep the viability of cells during normal growth as well as under MMS-induced replication-stressed conditions. On the contrary, cells without Ctf18 (0.003%) and Sgs1 (0.0015%) showed higher MMS-sensitivity than *mrc1Δ*, suggesting a role for these proteins in MMS-induced replication stress; and their corresponding *elg1Δ*-*DRC* mediator double mutants *elg1Δ ctf18Δ* and *elg1Δ sgs1Δ* showed a similar MMS-sensitivity to that of *ctf18Δ* and *sgs1Δ*, respectively (Fig. 1, G and H).

The observed synergistic relationship between Elg1 and DDR sensors/mediators in response to MMS-induced replication stress suggests that Elg1 acts in the S-phase checkpoint in parallel to the established sensors and mediators. Indeed, double mutants of DDR sensors (*ddc1Δ mec3Δ*, *mec3Δ rad17Δ*, *rad17Δ ddc1Δ* and *rad24Δ ddc1Δ*) and DDR sensors-mediators (*ddc1Δ rad9Δ*, *rad24Δ rad9Δ*, *rad24Δ mrc1Δ* and *ddc1Δ mrc1Δ*) do not show any additional MMS sensitivity compared to their respective single mutants (Fig. S1, B-E) indicating that the DDR sensors and mediators work through the same pathway to neutralize the MMS-induced replication stress.

The apical/sensor kinase Mec1 bridges DDR sensors, DDC/DRC mediators and the effector kinase Rad53, and therefore acts as a fulcrum of the S-phase checkpoint (43,44). As expected, mutation of *MEC1* is epistatic to deletion of any of the S-phase checkpoint pathway components, in response to MMS (Fig. S1F). This finding immediately suggests that *MEC1* should be in a synergistic relationship with *ELG1*, and indeed, as expected, *mec1Δ sml1Δ elg1Δ* cells found to be 6– and 24-fold more MMS sensitive than *mec1Δ sml1Δ* and *elg1Δ* cells, respectively (Fig. 1, I and J).

In summary, our extensive genetic analysis clearly established that under MMS-induced replication-stressed conditions, Elg1 functions in parallel to the DDR sensor-sensor kinase-mediator pathway.

### Elg1’s role in the DDR is PCNA-dependent

In the absence of Elg1, PCNA accumulates on DNA, and cells with accumulated chromatin-bound PCNA exhibit a number of defects in DNA/chromatin transactions such as increased sister chromatid recombination, elongated telomeres, inefficient DSB and mismatch repairs, defective nucleosome organization and inefficient chromatin silencing all of which lead to genome instability (7,13,16–18,45,46). The hypersensitivity of *elg1Δ*-*DDR* double mutants towards MMS compared to their corresponding single mutants revealed a replication stress-dependent synthetic genetic interaction between *ELG1* and *DDR* genes implying that Elg1 and S-phase checkpoint components collaboratively neutralize MMS-caused replication stress. However, the mechanism through which Elg1 relieves the replication stress is not known. To test whether PCNA retention on chromatin makes *elg1Δ*-*DDR* double mutants susceptible to MMS, we used a well-characterized PCNA disassembly-prone mutant (*pol30-D150E*) in which PCNA monomers, even in the absence of Elg1, fail to form a stable PCNA trimer ring on chromatin as the interaction between each monomer is disrupted due to a point mutation at the trimer interface (D150E) (13). If chromatin-bound accumulated PCNA is the reason for the high MMS-sensitivity of *elg1Δ*-*DDR* double mutants, then the same set of mutants in a *pol30-D150E* background (*pol30-D150E elg1Δ*-*DDR*) should show single *DDR* mutants-like sensitivity. Figure 2 shows that indeed this is the case: the sensitivity of all the *elg1Δ-DDR* sensor mutants (*elg1Δ ddc1Δ*, *elg1Δ mec3Δ*, *elg1Δ rad17Δ* and *elg1Δ rad24Δ*), as well as that of the *elg1Δ*-*DDC* mediator mutant (*elg1Δ rad9Δ*) was reduced to that of the single *DDR* mutants (Fig. 2A, C, E and G).

**Figure 2.**
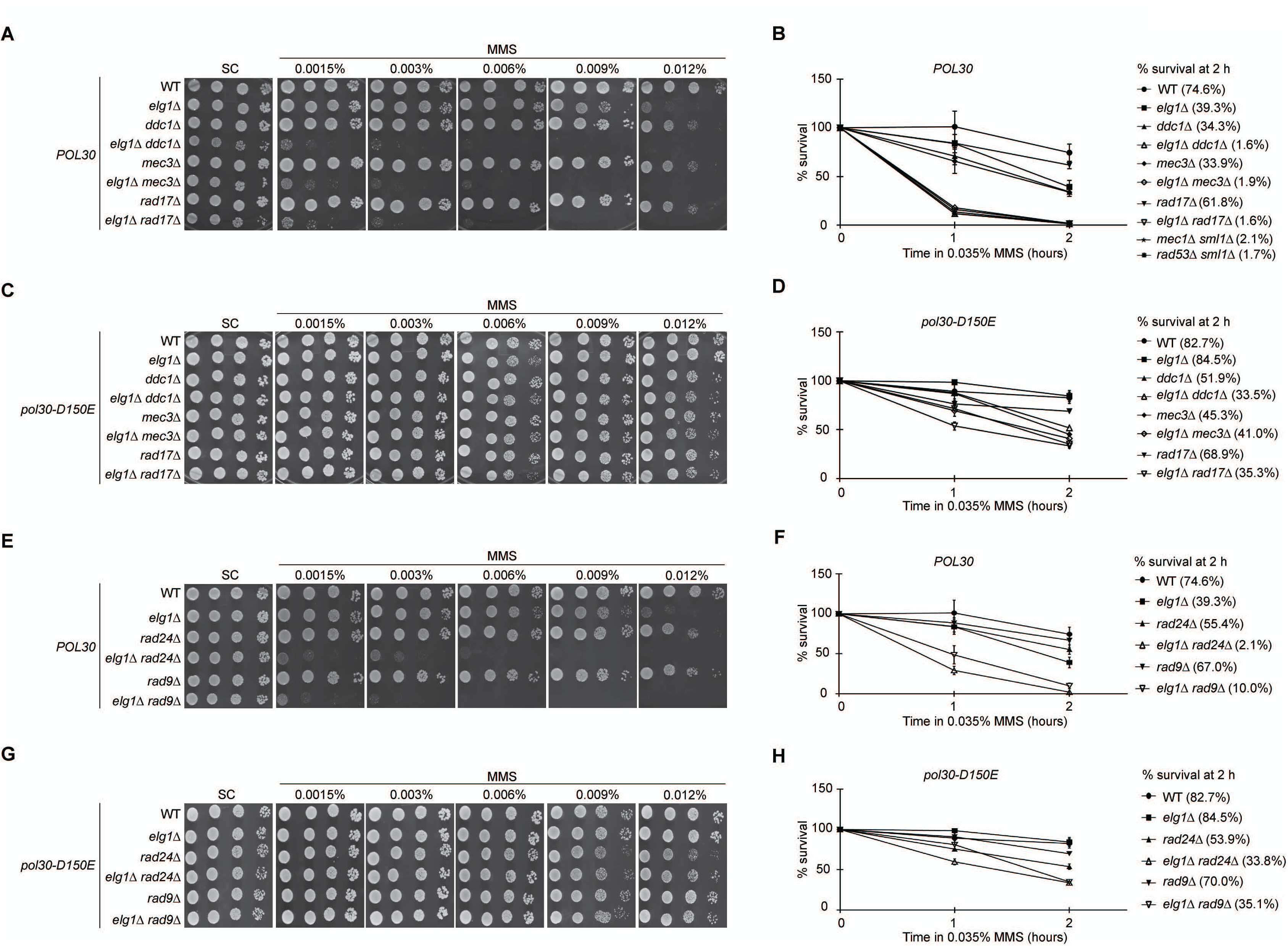
Disassembly-prone allele of PCNA restores the viability of MMS-treated *elg1Δ*-*DDR* mutants. (A) Ten-fold serial dilutions of overnight cultures of WT, *elg1Δ*, single mutants of 9-1-1 complex, and double mutants combining *elg1Δ* with 9-1-1 single mutants were dropped on SC plates supplemented with or without indicated amounts of MMS. (B) For the MMS survival assay of panel A strains as well as *mec1Δ sml1Δ* and *rad53Δ sml1Δ* mutants, exponentially growing cells at 28°C were exposed to 0.035% MMS for the indicated time points, and following sodium thiosulphate quenching, 10^2^-10^3^ cells were plated and incubated at 28°C. Survival was determined by counting the colony numbers of each time point, and the percentage was calculated in respect to 0 h colony number. (C) Drop assay of *pol30*-*D150E* (disassembly-prone PCNA mutant) panel A strains on indicated MMS containing SC plates, and (D) MMS survival assay of panel C strains. (E) Drop assay of WT, *elg1Δ*, *rad24Δ*, *rad9Δ* and double mutants coupling *elg1Δ* with *rad24Δ* and *rad9Δ* on indicated MMS containing SC plates, and (F) their MMS survival assay. (G) Drop assay of *pol30*-*D150E* panel E strains on SC plates supplemented with indicated amounts of MMS, and (H) their MMS survival assay. In MMS survival assay histograms, each data point is an average of three independent experiments, and the survival rates of the experimental strains, post-2 h of MMS exposure, are given. The error bars indicate ± SEM (standard errors of mean).

To further validate the drop assay result, we conducted cell survival assay where strains of both *POL30* and *pol30-D150E* background cells were exposed to 0.035% MMS for either one or two hours, following which cell viability was determined by scoring the number of colonies on rich nutrient plates (for details see Materials and Methods). In the *POL30* background, after 2 h of MMS exposure, the viability of WT and *elg1Δ* was found to be ∼75% and ∼39%, respectively whereas single *DDR* sensor mutants *ddc1Δ* and *mec3Δ* showed ∼34% viability, and *rad17Δ* and *rad24Δ* showed ∼62% and ∼55% viability, respectively (Fig. 2B and F). In contrast to single mutants, the percentage of viable cells after 2 h of MMS treatment for *elg1Δ*-*DDR* sensor double mutants (*elg1Δ ddc1Δ*, *elg1Δ mec3Δ*, *elg1Δ rad17Δ* and *elg1Δ rad24Δ*) dropped significantly to ∼2%, a value similar to the viability of 2 h MMS-treated *mec1Δ sml1Δ* and *rad53Δ sml1Δ* cells (henceforth *mec1Δ* and *rad53Δ* mutants), and hence displayed a synergistic loss of viability when *elg1Δ* was coupled with single *DDR* sensor mutants (Fig. 2B and F). When PCNA was blocked from accumulating on DNA (*pol30-D150E* background), the viability of *elg1Δ*-*DDR* sensor double mutants after 2 h of MMS exposure shot up from ∼2% (*POL30*) to ∼34-41% (*pol30-D150E*), similar to that of *POL30 DDR* sensor single mutants levels [compare Fig. 2B and D with Fig. 2F and H (for *elg1Δ rad24Δ*)]. The response of *elg1Δ*-*DDC* mediator double mutant (*elg1Δ rad9Δ*) to the MMS-exposed viability assay was similar to that of *elg1Δ*-*DDR* sensor double mutants as *POL30 elg1Δ rad9Δ* (∼10%) could barely grow whereas *elg1Δ rad9Δ* in *pol30-D150E* background (∼35%) restored its viability (Fig. 2F and H). In summary, the viability data not only upholds the drop assay results but also provides evidence that Elg1 employs its PCNA unloading activity to play an active role in the S-phase checkpoint following MMS-induced replication-stressed conditions.

### Passaging of *elg1Δ*-*DDR* mutants through S-phase with damaged DNA is lethal

Drop assay data showed that *elg1Δ*-*DDR* mutants are more sensitive to MMS than *mec1Δ sml1Δ* (Fig. 1I vs. Fig. 2A and E; Fig. S2) and *rad53Δ sml1Δ* (Fig. S2). Moreover, when logarithmically growing cells were exposed to a low dose of MMS, the cell viability of *elg1Δ*-*DDR* sensor double mutants (*elg1Δ ddc1Δ*, *elg1Δ mec3Δ* and *elg1Δ rad17Δ*) at 1 h (12 – 18%) and 2 h (∼2%) was found to be similar to that of *mec1Δ sml1Δ* cells (∼16% and ∼2% at 1h and 2h, respectively) and *rad53Δ sml1Δ* cells (∼12% and ∼2% at 1h and 2h, respectively) (Fig. 2B). The viability of *elg1Δ rad24Δ* at 1 h was slightly higher, but at 2 h it dropped to ∼2% equaling to the other *elg1Δ*-*DDR* sensor as well as to *mec1Δ* and *rad53Δ* mutants (Fig. 2F). By manipulating the timing of MMS-induced DNA damage it was found that *mec1Δ* and *rad53Δ* checkpoint mutants are extremely sensitive to MMS at S-phase compared to other phases, and traversing S-phase with damaged DNA results in extreme cytotoxicity of MMS in these checkpoint mutants (42). It turns out that in the presence of low dose of MMS, *mec1Δ* and *rad53Δ* checkpoint mutants are not able to complete chromosome replication as replication forks in these mutant cells arrest irreversibly, resulting in cell death (42,47). The similar survival rate of *elg1Δ*-*DDR* sensor and *mec1Δ*/*rad53Δ* mutants following MMS exposure during their exponential growth phase indicates that (like central checkpoint mutants) most likely the passaging of *elg1Δ*-*DDR* sensor double mutants through S-phase in the presence of MMS, is the reason of cell death. To test this, along with *mec1Δ* and *rad53Δ* mutants, *elg1Δ ddc1Δ* (a representative of *elg1Δ 9-1-1Δ* mutants) and *elg1Δ rad24Δ* cells were blocked at G1, and post-G1 release, cells were treated with 0.035% MMS for 1 h following which cells were plated for survival rate assessment. The viability of control checkpoint mutants *mec1Δ* (1.5%) and *rad53Δ* (1.9%) matched the previously published reports, whereas both *elg1Δ*-*DDR* sensor double mutants were found to be more sensitive to MMS (0.4% and 0.9% viability for *elg1Δ rad24Δ* and *elg1Δ ddc1Δ* mutants, respectively) compared to central checkpoint mutants [Fig. 3A; (42)]. Logarithmically grown *elg1Δ rad9Δ* cells showed 10% viability following a 2 h 0.035% MMS exposure; however, post-G1 synchronization, when these cells were exposed to 0.035% MMS for 1 h, their viability (1.5%) matched that of central checkpoint mutants (Figs. 2F and 3A). Together, this data confirms that in the presence of MMS, *elg1Δ*-*DDR* double mutants, like central checkpoint mutants, face cell death for passaging through S-phase with damaged DNA, and therefore, suggests that a combined absence of Elg1 and any one of the DDR sensor proteins or DDR mediator protein impedes chromosome replication, plausibly either by arresting forks irreversibly like *mec1Δ*/*rad53Δ* mutants or by blocking the functional pathways that resume DNA synthesis following stalling of replication forks.

**Figure 3.**
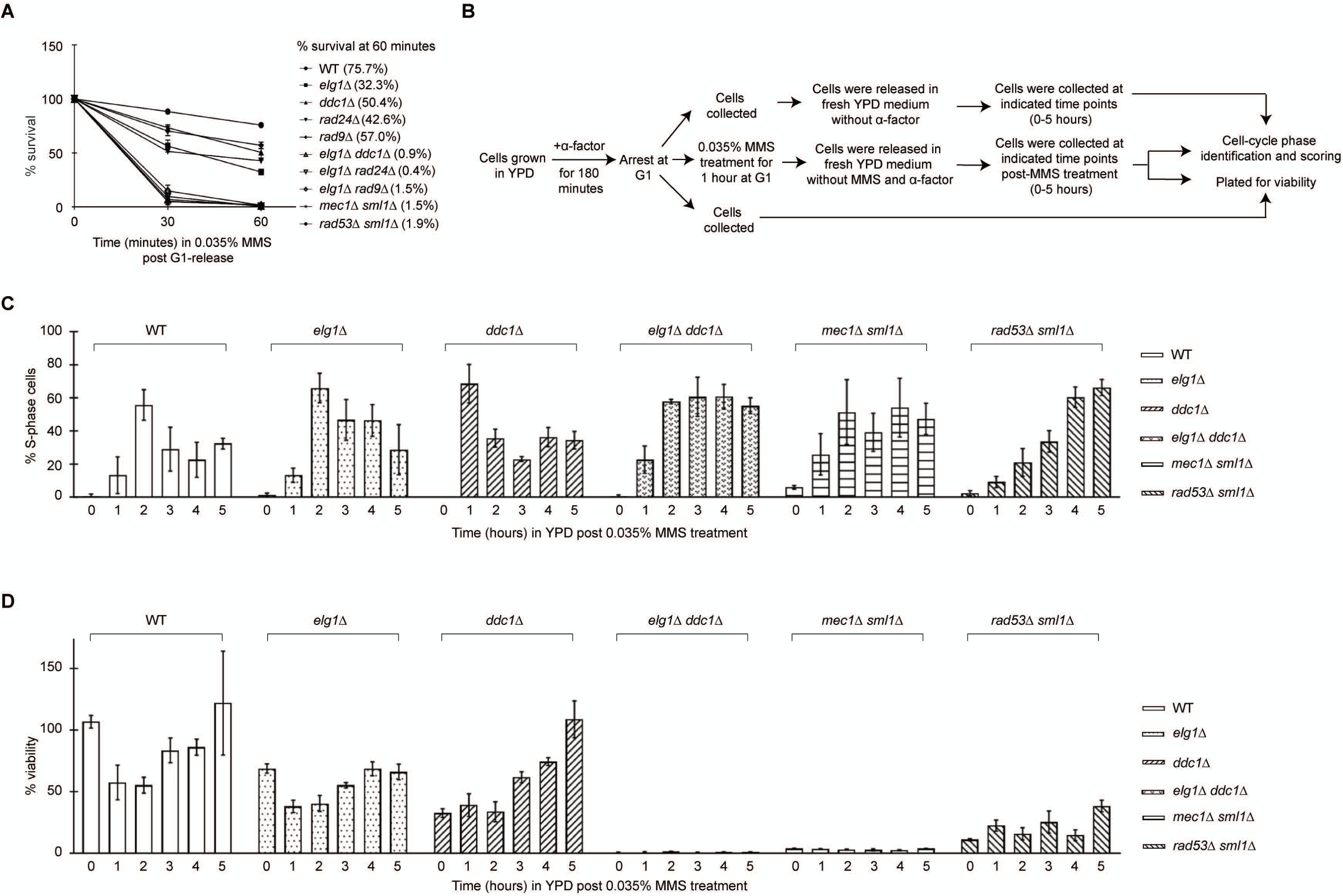
*elg1Δ ddc1Δ* cells, a representative of the *elg1Δ*-*DDR* mutants, fail to passage through S-phase with damaged DNA. (A) Following alpha factor-mediated G1 arrest, WT cells, central checkpoint mutants (*mec1Δ sml1Δ* and *rad53Δ sml1Δ*), selected *elg1Δ*-*DDR* double mutants (*elg1Δ ddc1Δ*, *elg1Δ rad24Δ* and *elg1Δ rad9Δ*) and their corresponding single mutants (*elg1Δ*, *ddc1Δ*, *rad24Δ* and *rad9Δ*) were released into 0.035% MMS-containing media. At 30 and 60 minutes, MMS-treated cells were aliquoted and following three-time ice-cold water wash, 10^2^-10^3^ cells were plated on rich nutrient-containing agar plates and incubated at 28°C for 3-4 days for viability scoring. Survival rates of the indicated strains at their aliquoted time points are displayed. Averages of three independent experiments were plotted. The error bars indicate ± SEM. (B) The schematic diagram represents the experimental regime that was employed to generate C and D. A set of G1-arrested cells (as indicated in C and D) were treated with or without 0.035% MMS for 1 h and then after releasing them into fresh media lacking either alpha factor or alpha factor and MMS, cells were collected in every 1 h interval to determine their cell cycle stages and cell viability. (C and D) The fraction of S-phase (small-budded cells) cells (C) and viable cells (D) present in WT, *elg1Δ*, *ddc1Δ*, *elg1Δ ddc1Δ*, *mec1Δ sml1Δ* and *rad53Δ sml1Δ* were scored at the indicated time points following a one-hour 0.035% MMS treatment in G1-arrested condition are displayed as bar graphs. Mean values for each time point and each cell-type were obtained by conducting three independent experiments. The error bars indicate ± SEM.

The cell viability of central checkpoint mutants remains low if cells are treated with MMS at G1, and subsequently released into MMS-free medium for S-phase progression (42). *mec1Δ* and *rad53Δ* mutants that passage through S-phase, fail to maintain the stability of replication forks when they encounter alkylated DNA (28,42). Next, to understand whether *elg1Δ*-*DDR* double mutants show a similar phenotype, we examined the viability of *elg1Δ*-*DDR* mutants in the absence of MMS, but in the presence of alkylated DNA. We induced MMS-specific DNA damage in G1-arrested WT, single and double mutants (here *elg1Δ ddc1Δ* cells represents *elg1Δ*-*DDR* mutants as the viability data of S-phase MMS treated *elg1Δ ddc1Δ*, *elg1Δ rad24Δ and elg1Δ rad9Δ* mutants were almost the same) following which, cells were released into medium lacking MMS, and thereafter cell-cycle stages and viability were scored at indicated time points (Fig. 3B). We reasoned that in the presence of alkylated DNA, if cells are able to complete chromosome replication despite of slow S-phase progression due to either replication fork stability or repair and restart issues, then higher cell viability with longer S-phase would be seen. Alternatively, if replication forks arrest irreversibly or collapse or cannot repair and restart following fork stalling, then along with long S-phase confinement, the cell viability will be extremely low.

Without being exposed to MMS, at 1 h from post-G1 release, WT, single and *mec1Δ* mutant cells were almost equally distributed in S and G2/M phase, while *elg1Δ ddc1Δ* and *rad53Δ sml1Δ* mutant cells were predominantly at S-phase (Fig. S3A). Slow S-phase progression of *elg1Δ ddc1Δ* mutant, like *rad53Δ* mutant, possibly indicates that Elg1 and Ddc1 together, play a coordinated role during normal DNA replication. Following MMS exposure at G1, except the *ddc1Δ* mutant, all other experimental strains mostly (∼70-90%) remained arrested at G1 even after 1 h from G1 release [Fig. S3B; (48)]. In the same time-frame, being a G1/S checkpoint-deficient strain, ∼70% of *ddc1Δ* mutant cells reached S-phase, however in the absence of Elg1, *ddc1Δ* mutant cells remained arrested at G1 [Fig. 3C and Fig. S3B; (49)]. Two hours of release from MMS in G1, except for the *rad53Δ* mutant where only ∼21% of cells reached S-phase, >50% cells of the other strains reached S-phase (Fig. 3C). WT and *elg1Δ* cells resumed cell cycle progression following 3 h and 5 h of G1 (MMS) release, respectively; however, the majority (∼50-60%) of *elg1Δ ddc1Δ*, *mec1Δ* and *rad53Δ* mutant cells remain confined at S-phase till the end of the experiment (Fig. 3C).

Although *elg1Δ* mutant cells showed almost a similar S-phase confinement as *elg1Δ ddc1Δ*, *mec1Δ* and *rad53Δ* mutants (Fig. 3C), their viability remained largely unaffected (∼38-69% vs. ∼58-86% of WT; Fig. 3D). Similarly, the viability of *ddc1Δ* mutant cells reached WT level with the increase of post-G1 release time (Fig. 3D). However, *elg1Δ ddc1Δ* mutant cells were found to be less viable than *mec1Δ* and *rad53Δ* mutants when exposed to MMS during G1 arrest (0.6% vs. ∼4% and ∼11%, respectively; Fig. 3D, 0 h) as well as during 1-5 h post-G1 (MMS) release (0.9-1.4% vs. 2.5-3.9% and ∼15-39%, respectively; Fig. 3D). The increased S-phase accumulation and extreme low viability of *mec1Δ* and *rad53Δ* mutants are due to their inability to complete chromosome replication as it has been shown previously that replication forks arrest irreversibly in these mutants while they traverse S-phase with alkylated DNA (28,42). Moreover, *mec1Δ* mutants show more fork catastrophes and MMS-sensitivity in S phase than *rad53Δ* mutants (42), explaining why in our experiments, the viability of *rad53Δ* mutants is higher than that of *mec1Δ* mutants. While passaging through S-phase with alkylated DNA, the higher cell death of *elg1Δ ddc1Δ* mutant cells than *mec1Δ* and *rad53Δ* mutants despite having extended S-phase suggests that the large S-phase population of *elg1Δ ddc1Δ* mutants is not because these cells take longer time to complete chromosome replication due to slow progression of replication forks; instead, their replication forks must have collapsed either due to compromised repair and restart activities or arresting irreversibly, which led to the termination of chromosome replication, and hence, cell death.

### *elg1Δ*-*DDR* mutants exhibit high number of chromosome breaks following MMS-induced replication stress

Rad52 foci act as a S-phase molecular marker for double strand breaks (DSBs) and DSB-associated recombinational repair centers (50). To assay whether *mec1Δ* /*rad53Δ*-like viability loss of *elg1Δ*-*DDR* double mutants in the presence of MMS or alkylated DNA stemmed from replication-associated chromosomal breaks, we scored the number of Rad52-GFP foci containing small-budded cells after releasing G1-arrested cells into 0.035% MMS-containing medium. Sixty minutes after G1 release, ∼47-56% of S-phase *elg1Δ*-*DDR* double mutant cells showed Rad52 foci, whereas *elg1Δ* and single *DDR* mutants exhibited ∼35% and ∼27-35% of S-phase cells with Rad52 foci, respectively (Fig. 4, A and B). Under the same experimental regime, MMS-treated *mec1Δ* and *rad53Δ* mutants showed ∼78% and ∼64% small-budded cells with Rad52 foci, respectively. (Fig. 4B). Therefore, under MMS-induced replication stress, *elg1Δ*-*DDR* double mutants show an incidence of Rad52 foci markedly higher than the single mutants but comparable to that of *mec1Δ* and *rad53Δ* S-phase cells. These results explicitly indicate that Elg1 and DDR sensor/mediator proteins work conjointly to maintain the replication fork integrity during replication stress, and their absence mirrors *mec1Δ /rad53Δ*-like consequences where DSBs are formed at replication forks compromising the replication fork integrity and resulting high cell death.

**Figure 4.**
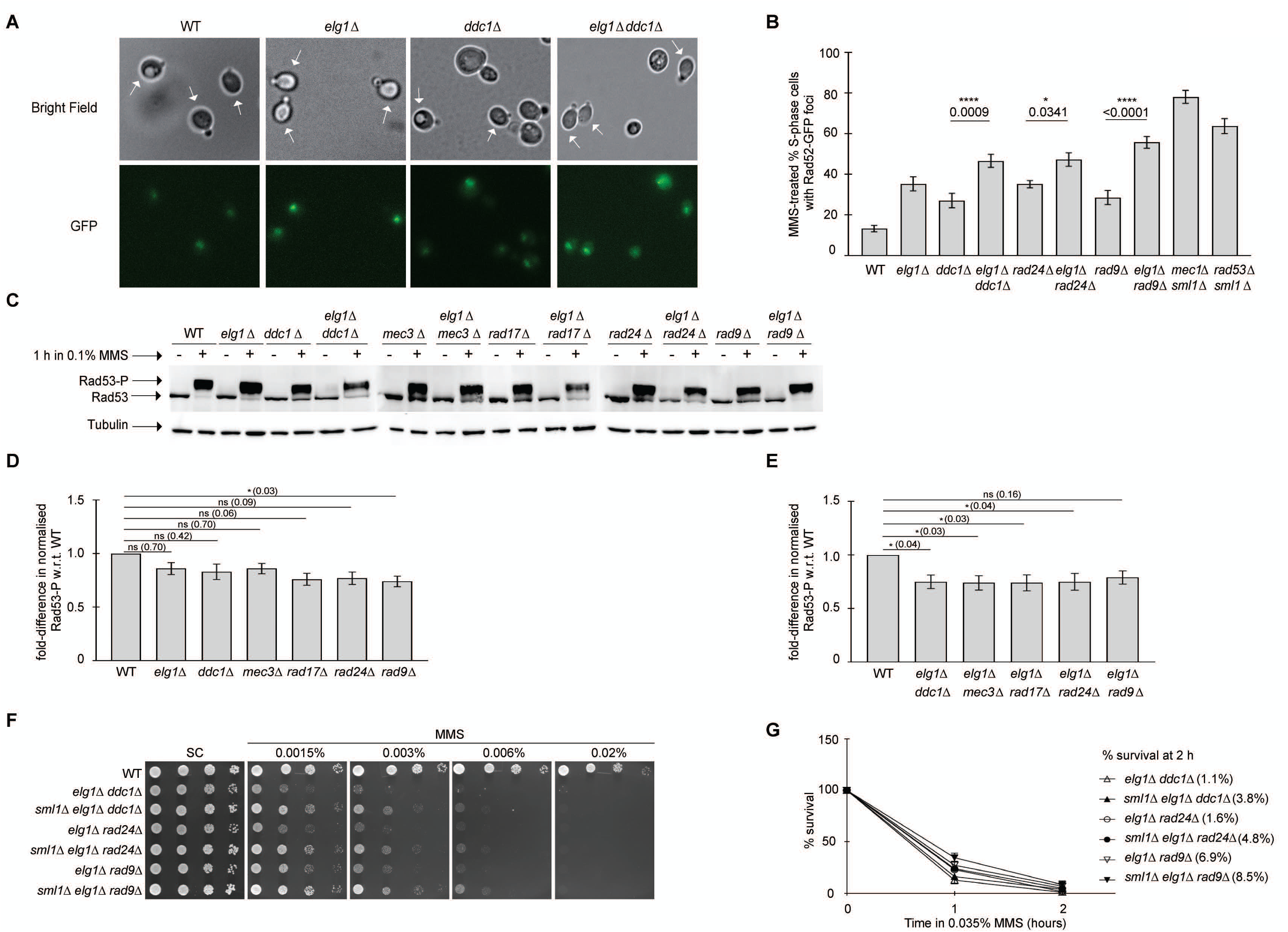
Under MMS-induced replication stress*, elg1Δ*-*DDR* mutants exhibit increased chromosome breaks but display a marginal compromise in Rad53 phosphorylation; and their lethality cannot be suppressed by elevating dNTP levels. (A-B) Exponentially grown WT and indicated mutant strains with Rad52-GFP were G1-arrested using alpha factor and then released into 0.035% MMS-containing medium for 60 minutes. Following the MMS treatment, only small-budded (S-phase) cells were analyzed for Rad52 foci scoring. (A) Representative cell images from the indicated strains are displayed. The S-phase cells are indicated with the arrow marks. (B) The percentage of S-phase cells with Rad52 foci for each experimental strain in response to MMS is displayed as a bar graph. Three sets of experiments were conducted and for each strain, combining all sets, at least 300 cells were counted. Data represent mean ± SEM. *P* values were obtained using one-way analysis of variance (ANOVA), and statistical significance tests of differences between two sets of mean values were done using Šídák’s multiple-comparison test. *P* value for each compared set is displayed in histograms. *, *P* < 0.01; *ns*, not significant. (C – E) Indicated strains were grown in YPD medium at 28°C till mid-log phase and then cells were exposed to 0.1% MMS for 1 h. Following genotoxic stress, proteins were extracted using trichloroacetic acid (TCA) method. Appropriate SDS-PAGE gels were run to resolve Rad53 and tubulin following which they were probed with appropriate antibodies as mentioned in the Materials and Methods section. (C) Level of Rad53 phosphorylation of indicated strains following 0.1% MMS exposure for 1 h. Tubulin was used as a loading control. (D – E) The intensities of Rad53 phosphorylation and tubulin of panel C strains were measured using ImageJ software. To calculate the fold-difference in Rad53 phosphorylation for all single (D) and double mutants (E) with respect to (w.r.t) WT Rad53 phosphorylation, first, individual Rad53 phosphorylation value of all strains was normalized using the corresponding tubulin value, and then the mutants’ individual normalized Rad53 phosphorylation value was divided by the normalized WT Rad53 phosphorylation value. For C – E, total ten independent experiments were done to obtain the normalized mean Rad53 phosphorylation value for each strain. The error bars indicate ± SEM. *P* values were obtained using one-way ANOVA and Šídák’s multiple-comparison test. *P* value for each compared set is displayed in histograms. *, *P* < 0.01; *ns*, not significant. (F) Overnight cultures of WT and representative *elg1Δ*-*DDR* double mutants with or without *sml1Δ* deletion were dropped on SC plates containing indicated amount of MMS. (G) Exponentially grown panel F strains were administered with 0.035% MMS and after ice-cold water wash for three times, 10^2^-10^3^ cells were plated. Viability of strains at 1 h and 2 h following MMS exposure was scored by counting the colony numbers of each time point, and the percentage was calculated in respect to 0 h colony number. Data, derived from three independent experiments, were displayed as mean ± SEM.

### Rad53 phosphorylation is altered in *elg1Δ*-*DDR* mutants in response to MMS

The Rad53 kinase, upon replication stress, undergoes phosphorylation in either a Mec1-Rad9 (DDC branch) or a Mec1-Mrc1 (DRC branch) pathway, and plays a pivotal role in executing the two most important functions of the S-phase checkpoint: upregulation of dNTP pool and stabilization of stalled replication forks (25,35,38,51). Hence, Rad53 phosphorylation is absolutely critical for the viability of the replication-stressed cells, and is considered to be the readout for an active S-phase checkpoint (28,42). The significant loss of viability of *elg1Δ*-*DDR* double mutants in response to MMS could be an effect of a non-functional Mec1-Rad53 axis. Therefore, we decided to test the phosphorylation of Rad53 in *elg1Δ*-*DDR* double mutants. Upon exposure to 0.1% MMS, compared to WT, *elg1Δ*, *ddc1Δ* and *mec3Δ* mutants showed a slight reduction in Rad53 phosphorylation (14-17%), whereas for *rad24Δ* and *rad17Δ* mutants, the reduction was comparatively higher (23% and 24%, respectively) but not statistically significant (Fig. 4, C and D). Only in *rad9Δ* cells, the phosphorylation level of Rad53 was significantly reduced (26%) (Fig. 4, C and D). Although in comparison to WT, except *elg1Δ rad9Δ*, the Rad53 phosphorylation levels of all other *elg1Δ*-*DDR* double mutants were found to be reduced significantly (∼26%) (Fig. 4, C and E); however, it was not completely abolished as in central checkpoint mutants (36,52). Together, these results provide clear evidence that under MMS-induced replication stress, unlike central checkpoint mutants *mec1Δ*/*rad53Δ*, the Mec1-Rad53 axis in *elg1Δ*-*DDR* mutants remains largely functional, and such marginal reduction in Rad53 phosphorylation cannot be solely responsible for their intense cell death.

### Elevated dNTP pool cannot rescue *elg1Δ*-*DDR* mutants from MMS-induced replication stress

Stressed replication forks induce the Mec1-Rad53 axis, activating the Rad53 kinase (53). Partial execution of this kinase cascade could in principle result in a shortage of dNTPs. To elevate the dNTP level, activated Rad53 phosphorylates the downstream protein Dun1, which in turn upregulates the dNTP pool by leading to the degradation of Sml1, an inhibitor of the ribonucleotide reductase enzyme complex, and by inhibiting Crt1, the repressor of *RNR* genes (25,39). To examine whether scarcity of dNTP in *elg1Δ*-*DDR* double mutants due to an inactive Dun1-Sml1 pathway is responsible for their cell death, we checked the growth of these mutants in the *sml1Δ* background on MMS-containing plates. The absence of Sml1 rescued *elg1Δ*-*DDR* double mutants only marginally in the presence of MMS (Fig. 4, F and G). *elg1Δ* exhibited an epistatic interaction with deletions of *RAD53* and *SML1* but showed an additive interaction with *dun1Δ* in response to MMS (Fig. S4). Together, these results rule out the possibility that upon MMS-induced replicative stress, the downstream activities following Rad53 phosphorylation are impaired in *elg1Δ*-*DDR* double mutants. Rather, these results confirm that in response to MMS-induced replication stress, the increased chromosome breaks at the replication forks of these double mutants (Fig. 4, A and B) are not due to the fork collapse mediated by dNTP shortage-dependent irreversible fork arrest, but possibly due to some other mechanisms such as replisome instability and/or inability in repairing/restarting damaged forks.

## Discussion

Although Elg1 is known to participate in several genome stability maintenance activities ranging from replication and repair-associated events to telomere length homeostasis, its role in replication stress response remains unclear. To address it, we combined *ELG1* deletions with DDR-compromised cells to examine the survival fitness of the derived *elg1Δ*-*DDR* double mutants under MMS-induced replication stress. The principal findings from the present study, summarized in Fig. 5, are (a) Elg1 counters MMS-induced replication stress through an independent pathway that works in parallel to that of the canonical S-phase checkpoint pathway. (b) The S-phase checkpoint role of Elg1 in rescuing DDR-compromised cells from MMS-induced replication stress manifests itself through unloading the chromatin-bound PCNA. (c) Despite the largely functional Mec1-Rad53 axis, *elg1Δ*-*DDR* double mutants, like central checkpoint mutants, fail to complete chromosome replication in the presence of alkylated DNA and display *mec1Δ*/*rad53Δ*-like mortality. And (d) Replication-stressed *elg1Δ*-*DDR* double mutants remain non-responsive towards increased cellular dNTP concentration, and show extensive dsDNA breaks, most likely, reflecting replication fork collapse. Collectively, our findings provide evidence that Elg1, in collaboration with canonical DDR sensors/mediators, uses a Mec1/Rad53-indepndent noncanonical pathway to counter MMS-induced replication stress, and thereby protects the genomic integrity of cells.

**Figure 5.**
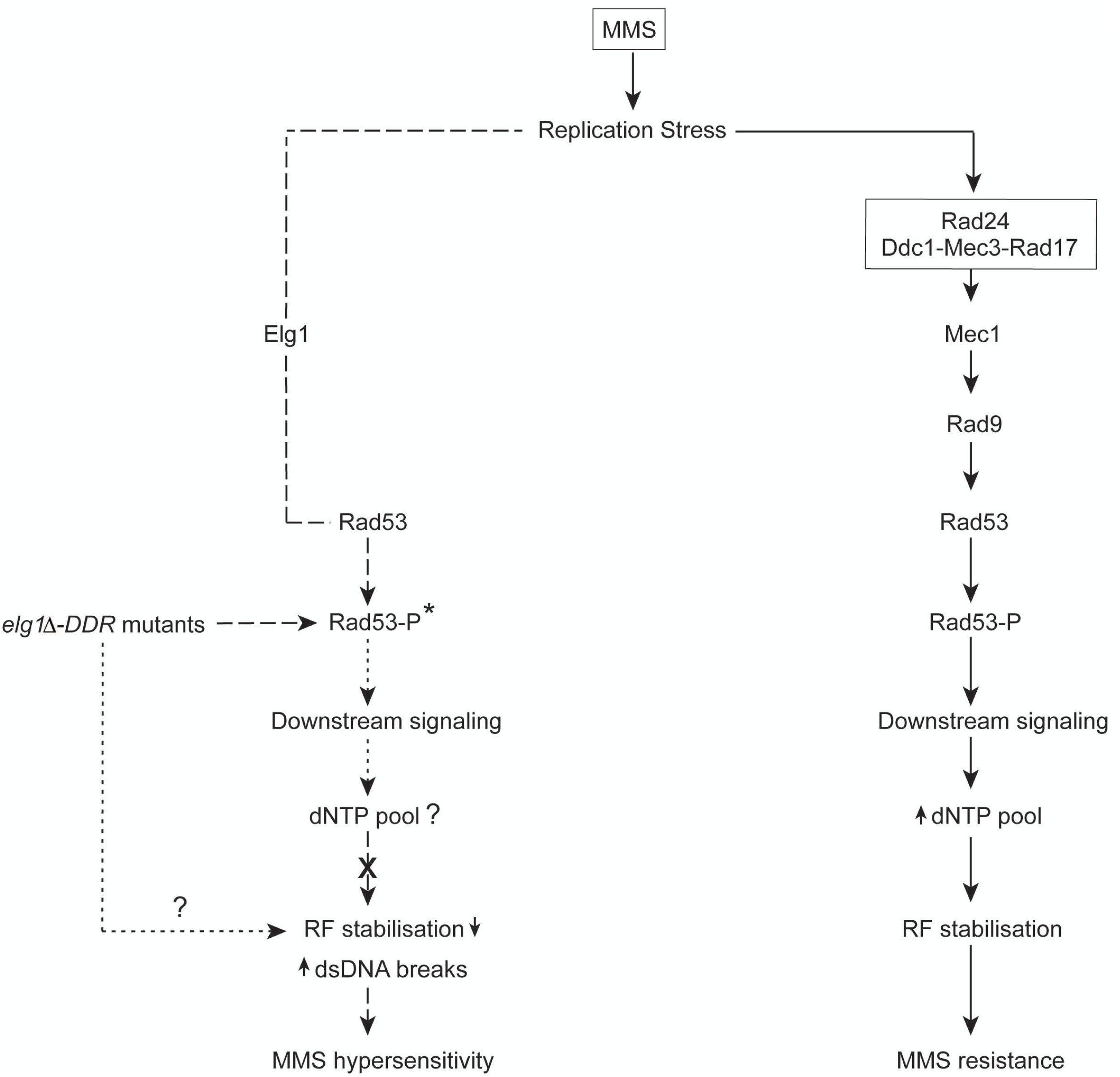
Counterpoising replication stress by Elg1-RLC: A model. A model embodying the current results displays a role for Elg1 in the S-phase checkpoint regulation as revealed by examining the *elg1Δ*-*DDR* double mutants. Genetic data sets uncover a completely PCNA-dependent Elg1-led pathway that runs in parallel to the established canonical S-phase checkpoint pathway to protect the viability of single DDR-compromised cells during MMS-induced replication stress. Although the cell biology data reveal that the increased chromosome breaks, most likely at the damaged/stalled replication forks, could be a possible reason for the death of *elg1Δ*-*DDR* double mutants under replication stress; however, the combined genetic and biochemical data do not endorse it to be a Rad53-controlled dNTP level-mediated replication fork destabilization event (indicated by X sign). * means marginal compromise in Rad53 phosphorylation in response to MMS. Solid lines represent published information, whereas dashed lines indicate the findings of this study. The question marks and dotted lines are for information yet not known.

Our results show that *elg1Δ* exhibits synergistic sensitivity to MMS in combination with the mutants of DDR sensors (9-1-1 mutants and *rad24Δ*; Fig. 1, A-D) and mediators (*rad9Δ* and *mrc1Δ*; Fig. 1, E-H), as well as with the mutant of the apical kinase (*mec1Δ*; Fig. 1, I and J). The emerged data unequivocally supports the notion that Elg1 contributes to replication stress response through a unique pathway that runs in parallel to the canonical S-phase checkpoint pathway (Fig. 5). Is Elg1 important in countering MMS-induced replication stress? In response to MMS, DDR-compromised cells show almost wild type-like growth. The simplest interpretation would be that DDR-compromised cells are devoid of only one of its DDR sensors or mediator proteins, therefore the remaining DDR protein pool along with active Mec1-Rad53 axis support their robust growth even under replication stress. However, despite the active presence of DDR protein pool-aided Mec1-Rad53 axis, withdrawal of Elg1 from DDR-compromised cells proves to be fatal for them. This clearly suggests that alongside canonical DDR pathway components, Elg1 plays an important role in neutralizing replication stress to maintain the viability of DDR-compromised cells in challenging conditions.

How does Elg1 safeguard DDR-compromised cells from MMS-induced replication stress? During normal replication, compared to leading strands, PCNA trimers are enriched on lagging strands to support the synthesis of Okazaki fragments (54). However, under replication stress, as Elg1 unloads PCNA only from lagging strands, PCNA trimers become enriched on the leading strands (54). Therefore, cells lacking Elg1 eventually accumulate PCNA on both strands of slowed down/stalled forks (54). We speculated that Elg1, by unloading PCNA from stressed forks of DDR-compromised cells, upkeep the viability of these cells under replication stress. The phenotypes observed in the PCNA disassembly-prone mutant background under MMS-induced conditions supported our view (Fig. 2, C and G). Due to the point mutation at the trimer interface (D150E), PCNA monomers cannot form PCNA rings to encircle dsDNA, and as a result, the chromatin of *pol30-D150E* strains does not accumulate PCNA. Drop assays as well as viability assays, performed under MMS-induced but PCNA-free chromatin conditions, showed a complete growth revival of *elg1Δ*-*DDR* double mutants as their growth matched that of DDR-compromised cells.

Next, to understand the severity of cell lethality of *elg1Δ*-*DDR* double mutants under replication stress, we compared their response with that of the central checkpoint kinase mutants *mec1Δ* and *rad53Δ*. Surprisingly, asynchronously grown *elg1Δ*-*DDR* double mutants and *mec1Δ/rad53Δ* mutants showed a similar death profile when treated with low dose of MMS. When cells are challenged with MMS, Mec1 and Rad53 proteins are required to maintain the integrity of replication forks for the completion of ongoing replication process (28,42). Our finding immediately raises the question: does contemporary absence of any one of non-essential DDR components and Elg1 affect the replication of MMS-induced stressed cells? To test this possibility, replication forks of *elg1Δ ddc1Δ* and central checkpoint kinase mutant cells were asked to passage through S-phase by encountering alkylated DNA. Bud morphology data clearly shows that following G1 release, like *mec1Δ* and *rad53Δ* mutants, the majority of the *elg1Δ ddc1Δ* cells are retained in S-phase for a prolonged time, indicating either slowing down of fork movement or inability of cells to complete replication. The cell viability data, derived from these cells, confirms the latter interpretation, and surprisingly, the intensity of *elg1Δ ddc1Δ* cell death surpasses that of central checkpoint kinase mutants. An almost null viability of *elg1Δ ddc1Δ* cells seems to suggest that the fraction of *elg1Δ ddc1Δ* cells that manages to passage through S-phase, could not remain alive either because they proceed for mitosis with partially replicated chromosomes or fail to segregate fully replicated chromosomes into daughter cells. Our work evidently demonstrates that under MMS-induced replication stress, accumulated PCNA on replicating strands indeed slows down the replication process; however, it is certainly not the cause of extreme cell lethality of *elg1Δ ddc1Δ* cells as in *elg1Δ* cells, post G1 release, half of the population are retained in S-phase for a long period but eventually almost two-third cells remain viable. But in the absence of a functional 9-1-1 clamp (generated due to the absence of Ddc1), PCNA accumulation on leading and lagging strands proves to be fatal for the cell. Though currently the reason is unknown, this result implies that in response to MMS, stressed cells are not able to cope with the co-absence of Elg1 and functional 9-1-1 clamp or its loader or DDC mediator.

We speculate that like *mec1Δ* and *rad53Δ* mutants, the active forks of *elg1Δ ddc1Δ* cells might have collapsed irreversibly by encountering damaged DNA, and therefore, failed to complete chromosome replication. The synergistic increase of Rad52 foci in the S-phase cells of *elg1Δ*-*DDR* double mutants while carrying out replication in the presence of alkylated DNA explicitly supports our argument. Generally, cells, facing the MMS challenge, activate the intra S-phase checkpoint to phosphorylate Rad53 to preserve the structural integrity of slowed down/stalled replication forks, and thereby maintain the viability (28,42,55). By encountering alkylated DNA, as *elg1Δ*-*DDR* double mutants and *mec1Δ*/*rad53Δ* mutants display equal cell death, it raises a possibility that in the absence of functional 9-1-1 clamp/ its loader Rad24 / mediator Rad9, the accumulated PCNA rings on slow moving/arrested forks might have blocked the Rad53 phosphorylation in *elg1Δ*-*DDR* cells, and hence forks have collapsed leading to incomplete chromosome replication. Our assay finds that the intra S-checkpoint of *elg1Δ*-*DDR* double mutants is only marginally compromised if challenged with MMS. Therefore, our data rules out inactive Rad53 as a possibility for the intense death of MMS-treated *elg1Δ*-*DDR* double mutant cells. Next, we reasoned that if fork collapse in *elg1Δ*-*DDR* cells is a dNTP-deprived consequence due to the modest compromise in Rad53 phosphorylation, then an increase in dNTP amount should be able to restore the viability of these cells. However, *SML1* deletion fails to rescue MMS-stressed *elg1Δ*-*DDR* double mutants. This result unequivocally suggests that although in response to MMS, cell death of *elg1Δ*-*DDR* mutants equals that of cells lacking Mec1/Rad53, the mechanism(s) of replication fork collapse must be different for these two groups of mutant cells.

A previous study by Karras et al. (56) suggested a collaboration between the two similar clamps: PCNA and 9-1-1, in rescuing cells from MMS-induced DNA damage. Our results support their model, and suggest that the unloading of PCNA by the Elg1-RLC becomes essential for viability in the presence of cytotoxic agents when the DDR pathway is compromised. Collectively, our work indicates that timely unloading of PCNA by Elg1 from damaged or stalled DNA is extremely crucial for the ability of cells to survive genotoxic attacks, and provides a second line of defense for the genome. As the DDR pathways and Elg1 are conserved from yeast to humans and play a central role in the development of cancer, the study of a similar human *elg1Δ*-*DDR* model would be beneficial from the therapeutic perspective.

## Materials and Methods

### Yeast strains and Genomic manipulation

Yeast strains used in this study are listed in Table S1. The list includes strain background, relevant genotypes and appropriate references. All strains used in this study represent MK166 background except Western blot and cell biological analysis in which W303 and S288C background strains were used, respectively. Employing standard PCR-based gene deletion method, endogenous genes were deleted using antibiotic resistance cassettes (57) following which gene specific primers were used for their confirmation. To generate disassembly-prone PCNA mutants, a primer set (bordering ∼700 nt upstream and ∼350 nt downstream of PCNA encoding gene *POL30*), and a template DNA with glutamate (E)-coding triplet instead of aspartate (D) coding at the 450^th^ position of *POL30* were used. Homologous recombination-mediated integration of the yielded PCR products at the primer site-specific generated *pol30-D150E* strains, and *POL30* flanking *NHP6B* disruptor *LEU2* served as a selection marker (58). The generated PCNA mutants were confirmed by standard DNA sequencing method.

### Cell growth and G1 arrest

For all experiments, yeast cells were grown in rich nutrient (YPD) media at 28°C unless otherwise stated. For G1 arrest, cells with ∼0.5 OD_610_ were initially treated with 5 µg/ml α-factor (1mg/ml) for 45 minutes following which an additional 2.5-5 µg/ml α-factor was added to the growing medium for another 135 minutes.

### Drop Assay

5 µl of 10-fold serially diluted overnight yeast cultures grown in YPD medium at 28°C were spotted on freshly prepared synthetic complete (SC) plates supplemented with or without MMS. SC plates without genotoxic agents served as a natural growth reader for mutants. SC plates without and with MMS were kept at 28°C for 2 and 3 days, respectively, and images were recorded thereafter. For a particular MMS concentration, if 1:1000 diluted (third dilution) mutant cells showed negligible or no growth, that specific MMS concentration was registered as MMS sensitivity for that particular mutant strain.

### Cell Survival Assay

Exponentially growing cells, with or without arresting at G1, were treated with 0.035% MMS and aliquots were taken at the indicated time points. Following the treatment, MMS-treated cells were either quenched with equal volume of 10% sodium thiosulphate for 20 minutes or washed with ice-cold water for three times. A total of 10^2^-10^3^ cells were plated on YPD plates and incubated at 28°C. After 3-4 days of incubation, the viability percentage of each time point was calculated against the colony number obtained in 0 h.

### Budding Index

Cell cycle stages were identified and scored based on the qualitative assessment of the bud size. Cells with bud circumference up to ≤20% of the mother cell circumference were classified as S-phase, while those with >40% to that of the mother cell circumference were categorized as G2/M cells. Cells with bud sizes between ∼20-40% of the mother cell were excluded from the scoring due to ambiguity in distinguishing S-phase from G2/M-phase.

### Protein extraction and Western blotting

Proteins were extracted from yeast cells treated with or without 0.1% MMS using trichloroacetic acid (TCA) method as described previously (59). Rad53 and tubulin were resolved using 8% and 10% SDS-polyacrylamide gel, respectively. Detection of Rad53 and its phosphorylation was done by mouse anti-HA antibody (Invitrogen 26183 – clone 2-2.2.14; 1:2500), and tubulin was detected using rat anti-tubulin (alpha) antibody (BioRad MCA78G – clone YOL1/34; 1:1000). Goat anti-mouse horseradish peroxidase (HRP) (Advansta –R-05071-500; 1:10,000) and goat anti-rat HRP (Thermo Fisher Scientific – 31470; 1:2500) secondary antibodies were used.

### Microscopy

Exponentially growing cells were arrested at G1 phase using alpha factor (α-factor) and then released into 0.035% MMS-containing medium for 1 h. Aliquoted cells were washed three times with ice-cold water following which cell fixation was done as described previously (60). 3-4 µl cell suspension was used to prepare microscopic slides. Images were taken using a fluorescence microscope (Zeiss Axio Observer Colibri 5) with a 63x objective lens.

## Supporting information

Bose et al._Supplementary file

## Acknowledgments

We thank Prof. Martin Kupiec, Tel Aviv University, Israel for strains and Dr. Shubhra Majumder, Presidency University, India for the microscope facility. This work was supported by DBT, Govt. of India (grant BT/RLF/Re-entry 45/2016 awarded to S.S.), with partial support from DST-SERB, Govt. of India (grant CRG/2020/005559 awarded to S.S.). PB was supported by DST-SERB fellowship, Govt. of India (grant CRG/2020/005559).

## Author Contributions

PB performed experiments, analyzed and interpreted the data, and prepared figures for the manuscript. SS conceived the project, designed experiments, analyzed and interpreted the data, and wrote the manuscript.

## Notes

### Competing Interest Statement

The authors have declared no competing interest.

