## Supplementary material for "The Elg1 Replication Factor C-like complex safeguards cells from replication stress through a noncanonical pathway independent of the Mec1-Rad53 axis": Bose et al._Supplementary file

### Supplemental material

Figure S1

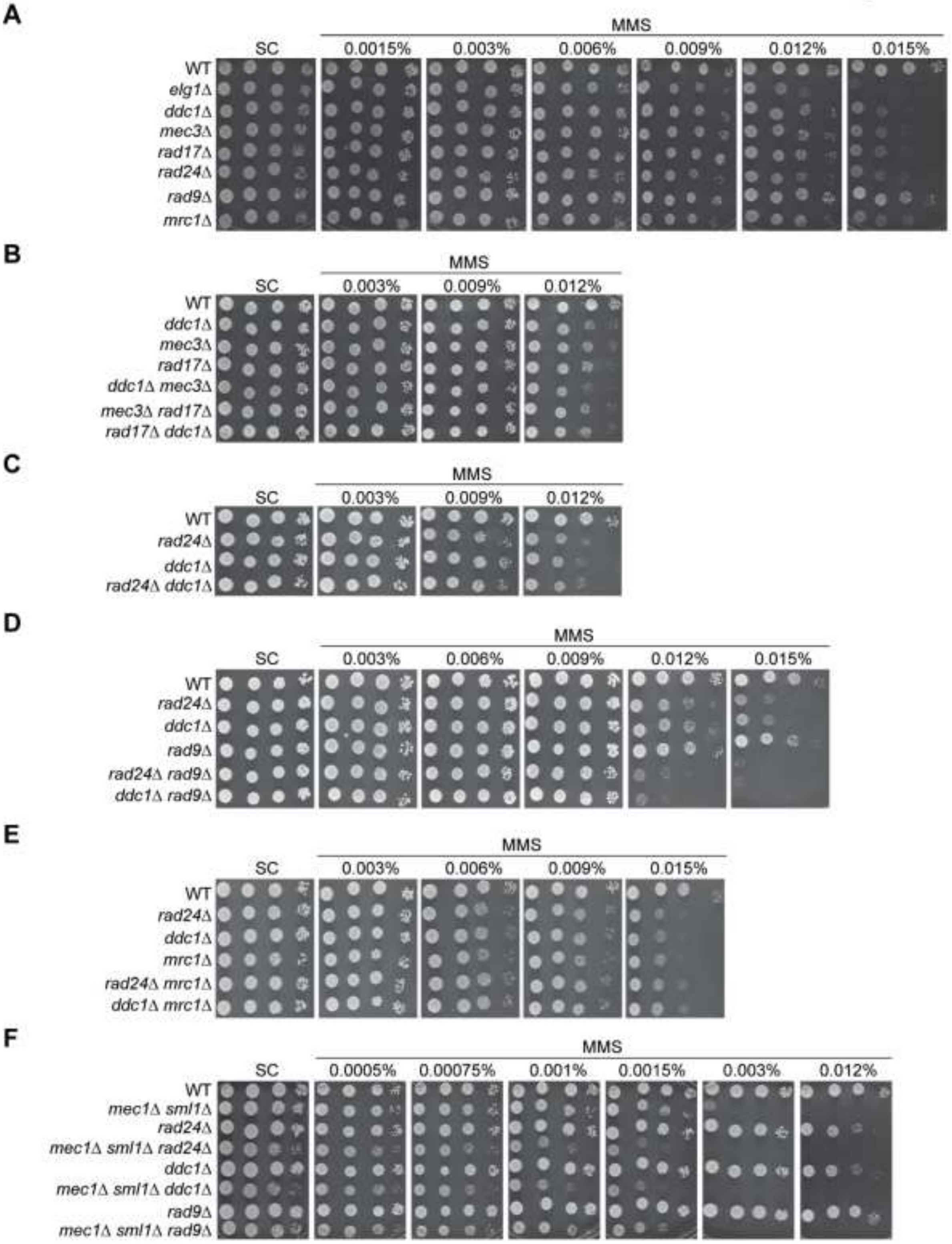

**Figure S1: Genetic interactions between different DNA Damage checkpoint components in response to MMS.** Ten-fold serial dilutions of overnight cultures of (A) *elg1Δ* and single *DDR* mutants, (B) single and double mutants of 9-1-1 complex, (C) *ddc1Δ*, *rad24Δ* and *rad24Δ ddc1Δ*, (D) *rad24Δ rad9Δ*, *ddc1Δ rad9Δ* and their corresponding single mutants, (E) *rad24Δ mrc1Δ*, *ddc1Δ mrc1Δ* and their corresponding single mutants, and (F) *mec1*Δ *sml1*Δ-*DDR* triple mutants and their corresponding single and double mutants were spotted on regular synthetic complete (SC) plates and SC plates containing indicated amounts of MMS. In each plate, WT spots were used as a control. For Figures S1, S2 and S4: for WT and *elg1Δ*, three technical repeats were carried out, whereas for other strains, three biological repeats were carried out. However, for the representation purpose, images from only one repeat is provided here. All SC plates ± MMS were kept at 28°C. Images of SC plates – MMS were recorded after 2 days of incubation, whereas SC plates + MMS were recorded after 3 days of incubation.

Figure S2

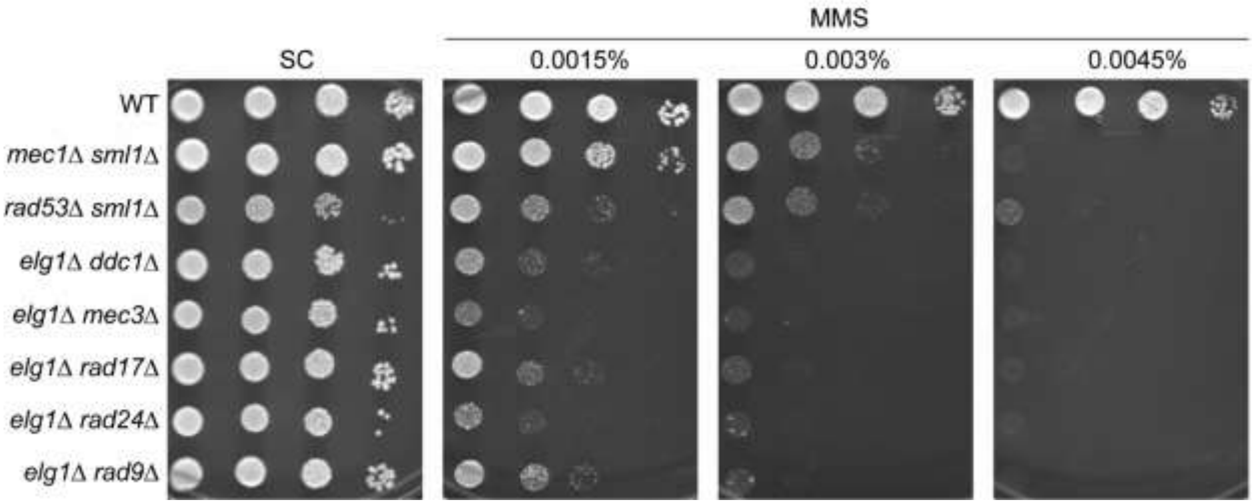


**Figure S2: *elg1Δ-DDR* mutants exhibit a marginally higher MMS sensitivity compared to the central checkpoint mutants *mec1Δ sml1Δ* and *rad53Δ sml1Δ*.** Overnight cultures of central checkpoint mutants and *elg1Δ-DDR* mutants were subjected to ten-fold serial dilution and spotted on SC plates supplemented with or without indicated amounts of MMS.

Figure S3

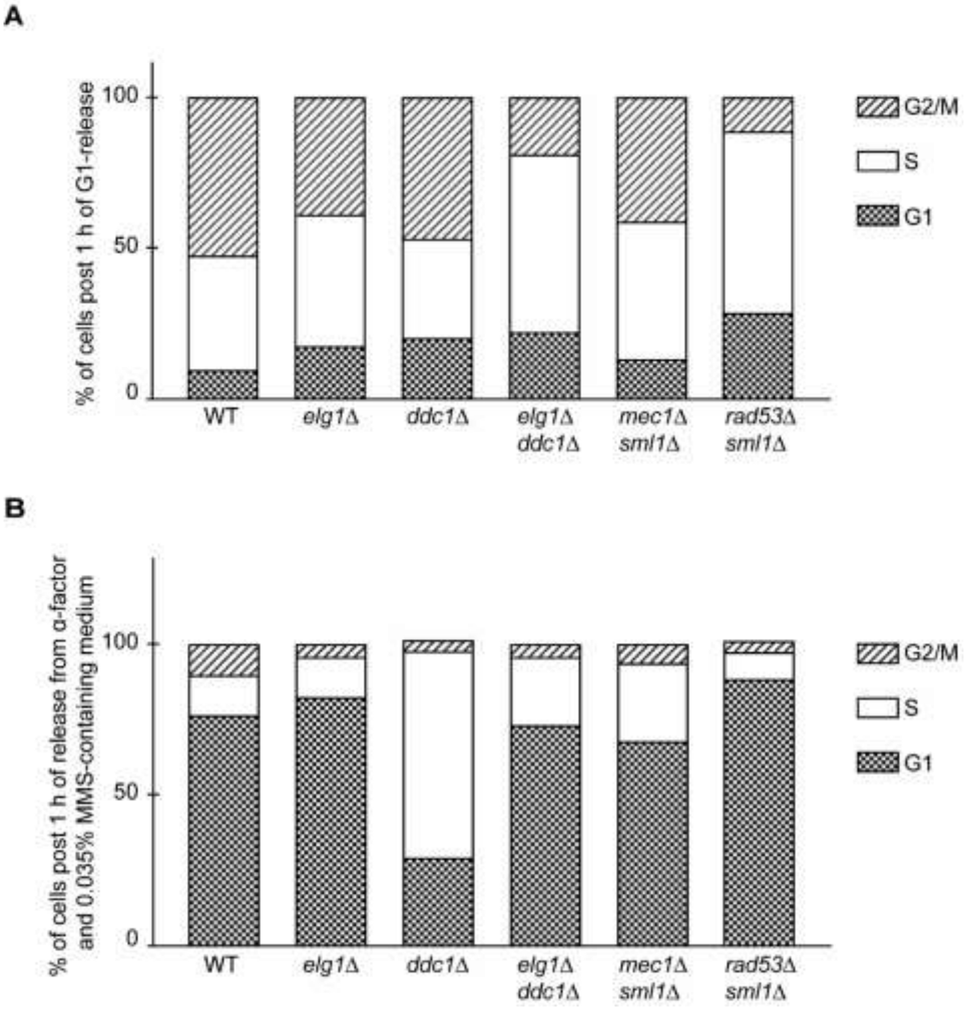


**Figure S3: Cell cycle distribution of WT and mutant cells following their release into MMS-free medium from (A) G1 arrest and (B) MMS-treated G1 arrest.** (A) WT and indicated mutants were arrested in G1 with α-factor and then released into MMS-free medium. Cells were collected after 1 hour and their cell-cycle stages were scored. Following G1 release, *elg1*Δ *ddc1*Δ displays a *rad53*Δ-like S-phase progression delay in MMS-lacking medium. (B) G1-arrested WT and indicated mutants were treated with 0.035% MMS for 1 hour and then released into fresh YPD medium. Cells were collected post 1 hour of YPD release and assessed for cell cycle stages. Except for *ddc1*Δ, the other indicated experimental cells mostly remained in G1 even after 60 minutes of release from MMS in G1.

Figure S4

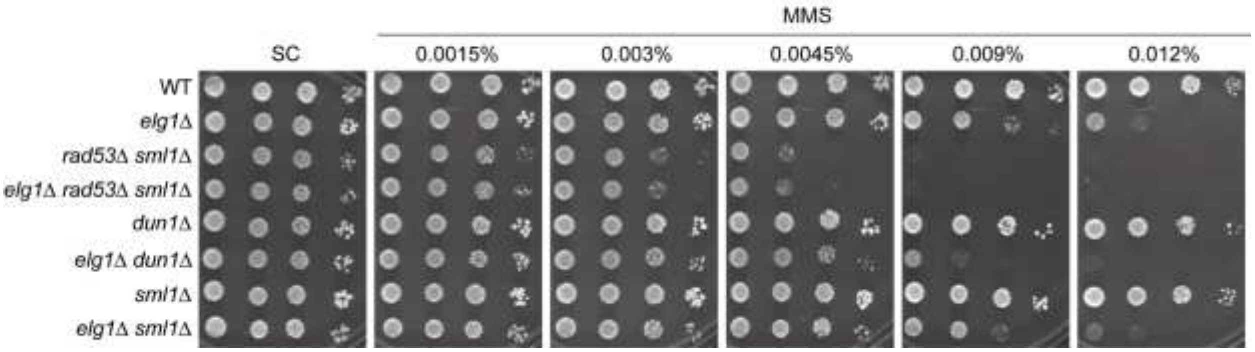


**Figure S4: In response to MMS, *elg1Δ* shows epistatic interaction with *sml1Δ* and *rad53Δ sml1Δ* but exhibits additive interaction with *dun1Δ*.** The genetic interactions of *ELG1* with *RAD53* / *DUN1* / *SML1* were examined under MMS-induced replicative stress by dropping ten-fold serially diluted overnight cultures of indicated single, double and triple mutants on SC plates containing indicated amounts of MMS.

**Table S1. Strains.** The yeast strains utilized for the present study are listed. The relevant genotypes of strains are indicated. The list is arranged roughly in accordance with the sequence in which the results are presented. An experiment in which a particular strain was utilized is denoted by qualifying it with the appropriate figure number. The relevant references are cited below.

| **Strain** | **Genotype** | **Relevant Figures** | **Source / Reference** |
|---|---|---|---|
| SSY19 Parent strain | MATa *lys2:: TySup ade2-1(0c) can1-100 (0c) ura3-52 leu2-3,112 trp1Δ901 HIS3:: lys2:: ura3- his4::TRP1::his4* | 1, 2, 3, 4, S1, S2, S3 and S4 | (Sau et al., 2019) |
| SSY6 Parent strain | SSY19 *Δelg1::HygMX*$^R$ | 1, 2, 3, S1, S3 and S4 | (Sau et al., 2019) |
| SSY587 | SSY19 *Δddc1::KanMX*$^R$ | 1, 2, 3, S1 and S3 | This study |
| SSY652 | SSY6 *Δddc1::KanMX*$^R$ | 1, 2, 3, 4, S2 and S3 | This study |
| SSY596 | SSY19 *Δmec3::KanMX*$^R$ | 1, 2 and S1 | This study |
| SSY656 | SSY6 *Δmec3::KanMX*$^R$ | 1, 2 and S2 | This study |
| SSY599 | SSY19 *Δrad17::KanMX*$^R$ | 1, 2 and S1 | This study |

| | | | |
|---|---|---|---|
| SSY659 | SSY6 *Δrad17::KanMX*$^{R}$ | 1, 2 and S2 | This study |
| SSY686 | SSY19 *Δrad24::KanMX*$^{R}$ | 1, 2, 3 and S1 | This study |
| SSY689 | SSY6 *Δrad24::KanMX*$^{R}$ | 1, 2, 3, 4 and S2 | This study |
| SSY666 | SSY19 *Δrad9::KanMX*$^{R}$ | 1, 2, 3 and S1 | This study |
| SSY669 | SSY6 *Δrad9::KanMX*$^{R}$ | 1, 2, 3, 4 and S2 | This study |
| SSY677 | SSY19 *Δmrc1::KanMX*$^{R}$ | 1 and S1 | This study |
| SSY314 | SSY19 *Δctf18::NatMX*$^{R}$ | 1 | This study |
| SSY917 | SSY19 *Δsgs1::HygMX*$^{R}$ | 1 | This study |
| SSY906 | SSY6 *Δmrc1::KanMX*$^{R}$ | 1 | This study |
| SSY316 | SSY6 *Δctf18::NatMX*$^{R}$ | 1 | This study |
| SSY907 | SSY6 *Δsgs1::KanMX*$^{R}$ | 1 | This study |
| SSY99 | MATa *lys2:: TySup ade2-1(0c) can1-100 (0c) ura3-52 leu2-3,112 trp1Δ901 HIS3:: lys2:: ura3-* | 1, 2, 3, S1, S2 and S3 | This study |

| | | | |
|---|---|---|---|
| | *his4::TRP1::his4 Δsml1::KanMX$^R$*<br>*Δmec1::HygMX$^R$* | | |
| SSY179 | SSY99 *Δelg1::URA3* | 1 | This study |
| SSY115 | SSY19 *Δsml1::KanMX$^R$*<br>*Δrad53::HygMX$^R$* | 2, 3, S2, S3 and S4 | This study |
| SSY360<br>Parent strain | RDKY8103 MATa<br>*pol30D150E::LEU2* | - | Gift from<br>Martin Kupeic |
| SSY1016 | SSY19 *pol30D150E::LEU2* | 2 | This study |
| SSY1019 | SSY6 *pol30D150E::LEU2* | 2 | This study |
| SSY1022 | SSY587 *pol30D150E::LEU2* | 2 | This study |
| SSY1037 | SSY652 *pol30D150E::LEU2* | 2 | This study |
| SSY1025 | SSY596 *pol30D150E::LEU2* | 2 | This study |
| SSY1040 | SSY656 *pol30D150E::LEU2* | 2 | This study |
| SSY1028 | SSY599 *pol30D150E::LEU2* | 2 | This study |

| | | | |
|---|---|---|---|
| SSY1043 | SSY659 *pol30D150E::LEU2* | 2 | This study |
| SSY1031 | SSY686 *pol30D150E::LEU2* | 2 | This study |
| SSY1034 | SSY689 *pol30D150E::LEU2* | 2 | This study |
| SSY1046 | SSY666 *pol30D150E::LEU2* | 2 | This study |
| SSY1049 | SSY669 *pol30D150E::LEU2* | 2 | This study |
| SSY528 | ATCC 201388: MATa *leu2Δ0 met15Δ0 ura3Δ0 Rad52-GFP::HISMX* | 4 | Gift from Martin Kupeic |
| SSY1462 | SSY528 *Δelg1::HygMX*$^{R}$ | 4 | This study |
| SSY1465 | SSY528 *Δddc1::KanMX*$^{R}$ | 4 | This study |
| SSY1468 | SSY1462 *Δddc1::KanMX*$^{R}$ | 4 | This study |
| SSY1472 | SSY528 *Δrad9::KanMX*$^{R}$ | 4 | This study |
| SSY1476 | SSY1462 *Δrad9::KanMX*$^{R}$ | 4 | This study |
| SSY1483 | SSY528 *Δrad24::KanMX*$^{R}$ | 4 | This study |

| | | | |
|---|---|---|---|
| SSY1487 | SSY1462 *Δrad24::KanMX*$^{R}$ | 4 | This study |
| SSY1493 | SSY528 *Δsml1::KanMX*$^{R}$ *Δmec1::HygMX*$^{R}$ | 4 | This study |
| SSY1498 | SSY528 *Δsml1::KanMX*$^{R}$ *Δrad53::HygMX*$^{R}$ | 4 | This study |
| SSY168 | W303 Mat a *ade2-1 Gal-Ddc2-LacI::HIS3 Rad53-HA::LEU2 LacO256::TRP1 GalS-Ddc1-LacI::URA3* | 4 | (Sau et al., 2019) |
| SSY171 | W303 Mat a *ade2-1 Gal-Ddc2-LacI::HIS3 Rad53-HA::LEU2 LacO256::TRP1 GalS-Ddc1-LacI::URA3 Δelg1::HygMX*$^{R}$ | 4 | (Sau et al., 2019) |
| SSY30 | W303 Mat a *ade2-1 Gal-Ddc2-LacI::HIS3 Rad53-HA::LEU2 LacO256::TRP1 GalS-Ddc1-LacI::URA3 ddc1Δ* | 4 | (Sau et al., 2019) |
| SSY123 | SSY30 *Δelg1::HygMX*$^{R}$ | 4 | (Sau et al., 2019) |
| SSY564 | SSY168 *Δrad17::KanMX*$^{R}$ | 4 | This study |
| SSY567 | SSY171 *Δrad17::KanMX*$^{R}$ | 4 | This study |

| | | | |
|---|---|---|---|
| SSY570 | SSY168 *Δmec3::KanMX$^R$* | 4 | This study |
| SSY1109 | SSY171 *Δmec3::KanMX$^R$* | 4 | This study |
| SSY1114 | SSY168 *Δrad24::KanMX$^R$* | 4 | This study |
| SSY1116 | SSY171 *Δrad24::KanMX$^R$* | 4 | This study |
| SSY1110 | SSY168 *Δrad9::KanMX$^R$* | 4 | This study |
| SSY1112 | SSY171 *Δrad9::KanMX$^R$* | 4 | This study |
| SSY386<br>Parent strain | SSY19 *Δsml1::NatMX$^R$* | - | This study |
| SSY1413 | SSY386 *(see above)*<br>*Δelg1::HygMX$^R$ Δddc1::KanMX$^R$* | 4 | This study |
| SSY1416 | SSY386 *(see above)*<br>*Δelg1::HygMX$^R$ Δrad24::KanMX$^R$* | 4 | This study |
| SSY1419 | SSY386 *(see above)*<br>*Δelg1::HygMX$^R$ Δrad9::KanMX$^R$* | 4 | This study |
| SSY594<br>Parent strain | SSY19 *Δmec3::KanMX$^R$* | - | This study |
| SSY708 | SSY594 *(see above)*<br>*Δddc1::HygMX$^R$* | S1 | This study |

| | | | |
|---|---|---|---|
| SSY597<br>Parent strain | SSY19 *Δrad17::KanMX$^R$* | - | This study |
| SSY704<br>Parent strain | SSY597 *(see above)*<br>*Δrad17::HygMX$^R$* | - | This study |
| SSY743 | SSY704 *(see above)*<br>*Δmec3::KanMX$^R$* | S1 | This study |
| SSY700 | SSY597 *(see above)*<br>*Δddc1::HygMX$^R$* | S1 | This study |
| SSY826 | SSY686 *Δddc1::HygMX$^R$* | S1 | This study |
| SSY688<br>Parent strain | SSY19 *Δrad24::KanMX$^R$* | - | This study |
| SSY939 | SSY688 *Δrad9::HygMX$^R$* | S1 | This study |
| SSY805 | SSY666 *Δddc1::HygMX$^R$* | S1 | This study |
| SSY1191 | SSY686 *Δmrc1::HygMX$^R$* | S1 | This study |
| SSY1195 | SSY587 *Δmrc1::HygMX$^R$* | S1 | This study |
| SSY1380 | SSY386 *Δmec1::HygMX$^R$*<br>*Δrad24::KanMX$^R$* | S1 | This study |
| SSY1377 | SSY386 *Δmec1::HygMX$^R$*<br>*Δddc1::KanMX$^R$* | S1 | This study |
| SSY1382 | SSY386 *Δmec1::HygMX$^R$*<br>*Δrad9::KanMX$^R$* | S1 | This study |
| SSY194 | SSY115 *(see above) Δelg1::URA3* | S4 | This study |

| | | | |
|---|---|---|---|
| SSY1383 | SSY19 *Δdun1::HygMX*$^R$ | S4 | This study |
| SSY1385 | SSY1383 *(see above)*<br><br>*Δelg1::URA3* | S4 | This study |
| SSY66 | SSY19 *Δsml1::KanMX*$^R$ | S4 | This study |
| SSY70 | MK166 MATα *Δsml1::KanMX*$^R$<br><br>*Δelg1::HygMX*$^R$ | S4 | This study |
